# miR-34c-3p regulates PKA activity independent of cAMP via ablation of PRKAR2B in *Theileria annulata*-infected leukocytes

**DOI:** 10.1101/2020.04.11.037341

**Authors:** Malak Haidar, Shahin Tajeri, Laurence Momeux, Tobias Mourier, Fathia Ben-Rached, Sara Mfarrej, Zineb Rchiad, Arnab Pain, Gordon Langsley

**Author notes:** Co-first authors (Malak Haidar, Shahin Tajeri).

## Abstract

MicroRNAs (miRNAs) are small non-coding RNAs that can play critical roles in regulating various cellular processes including during many parasitic infections. Here, we report a regulatory role for miR-34c-3p in cAMP-independent regulation of PKA activity in *Theileria annulata* infection of bovine leukocytes. We identified *prkar2b* (cAMP-dependent protein kinase A type II-beta regulatory subunit), as a novel miR-34c-3p target gene and demonstrated how infection-induced up-regulation of miR-34c-3p in leukocytes repressed PRKAR2B expression to increase PKA activity and promote the virulent disseminating tumour phenotype of *T. annulata*-transformed macrophages. Finally, we demonstrate that miR-34c-3p regulation of *prkar2b* expression is generalizable, by confirming that *Plasmodium falciparum* infection of red blood cells also raises intracellular levels of miR-34c-3p and show that this negatively regulates host *prkar2b* expression so increasing PKA activity. Infection-induced increase in miR-34c-3p levels, therefore, represents a novel cAMP-independent way of regulating host cell PKA activity in infections by *Theileria* and *Plasmodium* parasites.

**Abstract Importance:** *Theileria* and *Plasmodium* infections of leukocytes and erythrocytes; respectively, lead to an increase in host cell miR-34c-3p levels and we identified *prkar2b* (cAMP-dependent protein kinase A type II-beta regulatory subunit), as a specific miR-34c-3p target gene. We demonstrate how infection-induced up-regulation of miR-34c-3p repressed PRKAR2B expression to increase PKA activity independent of fluxes in cAMP. Thus, in two different host-parasite combinations infection-induced increase in miR-34c-3p represents a novel epigenetic way of increasing in host PKA activity that contributes to the pathology of disease.

## Introduction

MicroRNAs are a class of small non-coding RNAs and are key regulators in several biological processes ranging from development and metabolism to apoptosis and signalling pathways (1, 2). Indeed, their profiles are altered in many human diseases and particularly in cancer (3, 4), making them a major focus of research. The up-or down-regulation of miRNAs can cause drastic changes to gene expression due to the fact that miRNAs have up to hundreds of potential mRNA targets (5). Changes to gene expression can lead to malignant transformation, especially if miRNA targets include genes that regulate cell homeostasis, proliferation, cell cycle progression, adhesion, or dissemination (6).

Post-transcriptional control of gene expression by miRNAs is also increasingly recognized as a central part of host/pathogen interactions. The role of miRNAs in bacterial (7, 8), viral (9), parasitic helminth (10) and protozoan infections is now well established. *Plasmodium* and *Theileria* parasites are obligate intracellular protozoa of the phylum Apicomplexa. The goal for *Plasmodium* and *Theileria* parasites is to infect their invertebrate arthropod hosts (mosquitoes and ticks, respectively) for successful transmission and continuity of the life cycle with the sexual forms – thus ensuring exchanges of genetic material via genetic recombination. To ensure transmission to blood-sucking vectors the parasite population is amplified in the mammalian host. Post-inoculation by the insect vector, sporozoites of both species invade MHC class I- and class II-positive cells (hepatocytes for *Plasmodia* and leukocytes for *Theileria*), but thereafter their life cycles differ, as the *Theileria* parasite population amplifies within the infected leukocyte. Apicomplexan parasites are able to alter the global gene expression patterns of their respective host cells by interfering with signalling cascades to neutralize host defences (11) and improve their capacity to infect, proliferate and disseminate. Apicomplexa also manipulate their host cell’s miRNomes to their own benefit (12–14). For example, we have shown that miR-126-5p by directly targeting and suppressing JNK-Interacting-Protein JIP-2 liberates cytosolic JNK to translocate to the host cell nucleus and phosphorylate c-Jun that trans-activates AP-1-driven transcription of *mmp9* to promote tumour dissemination of *Theileria*-transformed macrophages (15). Moreover, miR-155 is also induced by infection to suppress De-Etiolated Homolog 1 expression that diminishes c-Jun ubiquitination (16). An increase in c-Jun levels leads to an augmentation in *BIC* (pri-miR-155) transcripts that contain miR-155, explaining how a positive feedback loop contributes to the growth and survival of *Theileria*-infected leukocytes (16). Furthermore, exosomes and their miRNA cargo play an important role in the manipulation of the host cell phenotype and the pathobiology of both *Theileria* and *Plasmodium* infections (17, 18). However, in contrast to *Toxoplasma,* the genomes of *T. annulata* and *P. falciparum* lack orthologs of Dicer and Argonaute, crucial enzymes in miRNA biogenesis (19, 20). Moreover, sequencing and bioinformatics analyses of small RNA libraries from *P. falciparum*-infected erythrocytes did not identify parasite-specific miRNAs (21). That said, at least 100 different human miRNAs are taken up by the parasite with a particular enrichment of miR-451 and let-7i in parasitized HbAS and HbSS erythrocytes (22). LaMonte *et al*. confirmed that human miRNA transferred into the parasite formed chimeric fusions with *P. falciparum* mRNA via impaired ribosomal loading, resulting in translational inhibition, eventually impairing parasite biology and survival. It is not yet known what determines the specific enrichment of particular miRNAs, or its incorporation into specific parasite mRNAs (20, 23). Extracellular Vesicles (EV) derived from *P. falciparum*-infected red blood cells (iRBC) contain miRNAs that can modulate target gene expression in recipient host cells and multiple miRNA species in EVs were identified bound to AGO2 forming functional complexes (18). Furthermore, *P. falciparum* takes up micro-vesicles containing AGO2 and miRNA from infected RBCs (24).

Herein, we demonstrate that by targeting *prkar2b* coding for the Protein Kinase A type II-beta regulatory subunit, miR-34c-3p regulates mammalian PKA activity independently of cAMP (Cyclic Adenosine Monophosphate) in *Theileria*-infected macrophages. We have previously shown that attenuated *T. annulata*-infected macrophages used as live vaccines against tropical theileriosis have diminished PKA activity and consequently impaired dissemination potential (25–27). We now show that attenuated macrophages have lower levels of miR-34c-3p and correspondingly higher levels of PRKAR2B that contribute to their diminished PKA activity. Significantly however, upon overexpression of miR-34c-3p in attenuated macrophages, both *prkar2b* transcripts and PRKAR2B protein levels drop leading to increased PKA activity that re-establishes the hyper-disseminating tumour-like phenotype of virulent macrophages.

Erythrocyte infection by *P. falciparum* also leads to an increase miR-34c-3p levels (18), so we asked if this led to an increase in infected erythrocyte PKA activity. We confirm that infection of red blood cells by *P. falciparum* leads to increased amounts of miR-34c-3p and demonstrate that inhibition of miR-34c-3p binding to its cognate seed sequence raised *prkar2b* transcript levels and reduced infected red blood cell PKA kinase activity. So, in two different host-parasite combinations infection-induced increases in miR-34c-3p lead to a reduction in *prkar2b* transcripts and an increase in host PKA activity independent of fluxes in cAMP.

## Results

### miR-34c-3p is differentially expressed in *T. annulata*-infected leukocytes

*T. annulata*-transformed virulent macrophages disseminate in infected animals causing a disease called tropical theileriosis. However, upon long-term in vitro passaging their capacity to disseminate becomes attenuated and they are used as live vaccines to control disease (21). We focused on miR-34c, as its expression is upregulated in *Theileria*-infected virulent macrophages, but is significantly down-regulated following attenuation of their dissemination capacity (Figure 1 left and right panel). We artificially increased miR-34c-3p levels of attenuated macrophages by transfecting them with a miR-34c-3p mimic (Fig 1B, left panel). Transfection of the mimic specifically increases miR-34c-3p levels, as by contrast endogenous miR-34c-3p levels did not change when attenuated macrophages were transfected with miR-30f (Fig. 1B, right panel).

**Figure 1:**
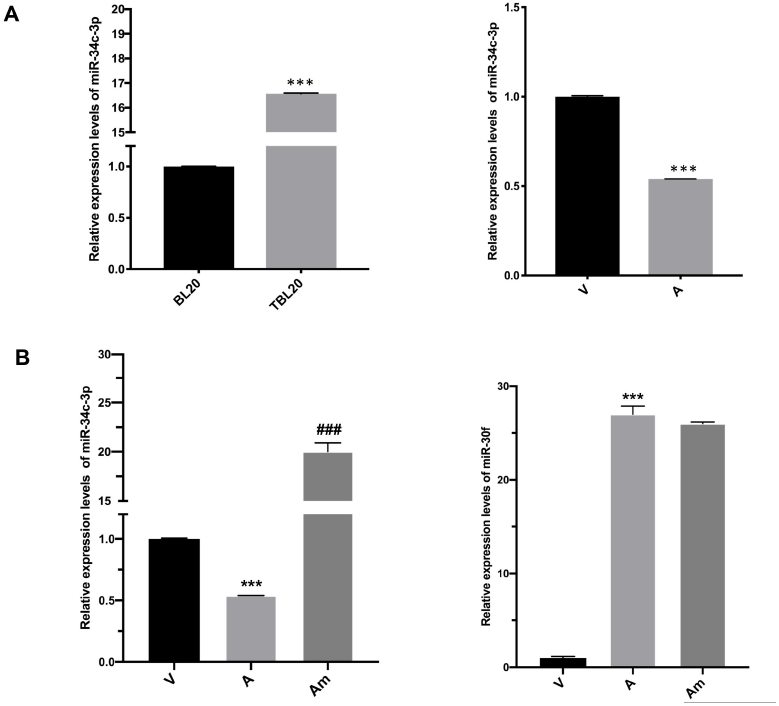
miR-34c-3p is differentially expressed in *T. annulata*-infected leukocytes. A left. miR-34c-3p levels in uninfected B lymphocytes (BL20) and levels increase when BL20 are infected with *T. annulata* (TBL20). **A right.** Levels of miR-34c-3p are high in virulent macrophages and decrease in attenuated macrophages. **B left panel**. miR-34c-3p levels in attenuated macrophages increase following their transfection with miR-34c-3p mimic. **B right panel**. Transfection of attenuated macrophages with the miR-34c-3p mimic specifically augments only miR-34c-3p levels, since amounts of miR-30f in attenuated macrophages are unaltered following transfection of the miR-34c-3p mimic. Data represented as mean ±SEM. *n*=3. \*\*\**P <* 0.001 compared to V; ###*P <* 0.001 compared to A. V stands for virulent, A for attenuated, Am: attenuated transfected with miR-34c mimic.

### Changes in miR-34c-3p levels influence the dissemination of *T. annulata*-transformed macrophages

Virulent macrophages display a high capacity to traverse Matrigel (22), and as miR-34c levels are upregulated in virulent macrophages its regulatory effect on the capacity of *T. annulata*-transformed macrophages to traverse Matrigel was examined. Transfection of attenuated macrophages with the miR-34c-3p mimic restored Matrigel traversal to levels equivalent to virulent macrophages (Figure 2). Thus, demonstrating that infection-induced increase in miR-34c-3p levels contributes to aggressive dissemination of *T. annulata*-transformed macrophages and that a drop in miR-34c-3p levels underpins the reduced dissemination of attenuated macrophages.

**Figure 2:**
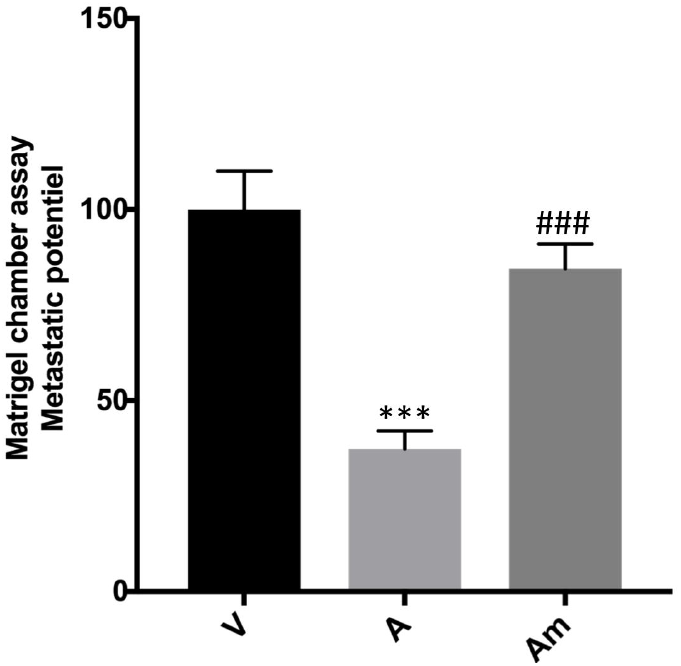
miR-34c-3p levels impact on dissemination potential of *T. annulata*-transformed macrophages. The ability of virulent macrophages (V) to migrate in Matrigel is higher compared to attenuated macrophages (A). Overexpression of miR-34c-3p in attenuated macrophages (Am) restores Matrigel traversal to virulent macrophage (V) levels. Data represent mean ±SEM. \*\*\**P<* 0.001 compared to V and ### *P*<0.001 compared to A.

### *prkar2b* is a direct target gene of miR-34c

miRNAs can regulate gene expression by binding to the 3’-untranslated region (3’UTR) of target mRNA. Since above we showed that miR-34c-3p expression can modulate the dissemination phenotype of *Theileria*-transformed macrophages, we looked for potential target genes that could explain the mechanism by which this is achieved. Using multiple algorithms (miRDB, miRWalk and TargetScan) *prkar2b* is predicted to be a miR-34c-target gene in both *Bos taurus* and *Homo sapiens*. In addition, we applied another criterion requiring that the predicted seeds occurred in genes, whose RNAseq-determined expression was low in virulent (where miR-34c levels are high) and high in attenuated (where miR-34c levels are low) (15). Combined this identified a canonical miR-34c-3p seed sequence in the 3’-UTR of *prkar2b*, one of the 4 genes coding for regulatory subunits of PKA.

Mammalian PKA is a hetero-tetrameric enzyme composed of two catalytic subunits associated with two regulatory subunits. The catalytic subunit PKA-C is bound to an inhibitory regulatory subunit (PKA-R), but upon binding of cAMP to the regulatory subunits, catalytic subunits are released to act as a serine/threonine kinase in both the cytoplasm and nucleus to phosphorylate the target proteins. PKA-C substrates include transcription factors and other proteins involved in developmental processes (30). We have previously shown that cAMP levels and PKA activity influence the dissemination potential of *Theileria*-transformed macrophages (25) and as *prkar2b* is one of four genes coding for regulatory subunits of PKA we determine if *prkar2b* is a bona fide miR-34c-3p target. First, *prkar2b* mRNA levels were measured in virulent and attenuated macrophages and following transfection of attenuated macrophages with the miR-34c-3p mimic (Figure 3A, left panel). Consistent with it being a miR-34c-3p target, *prkar2b* transcripts are more abundant in attenuated macrophages that have lower miR-34c-3p levels (see Figure 1) and PRKAR2B protein levels decreased in attenuated macrophages transfected with the miR-34c-3p mimic (Fig. 3A, right panel). To confirm that *prkar2b* is a direct target gene its 3’-UTR harbouring an identified canonical miR-34c-3p seed sequence was subcloned into the psiCHECK-2 (Promega, # C8021) and transfected into attenuated macrophages together with the miR-34c-3p mimic and Renilla luciferase activity monitored (Fig. 3B, left panel). Due to attenuated macrophages having lower levels of miR-34c-3p they display greater luciferase activity than virulent macrophages. Upon mimic treatment luciferase activity in attenuated macrophages dropped significantly confirming that *prkar2b* is a direct target of miR-34c-3p. In addition, we synthesized full-length *prkar2b* with potential miR-34 seeds mutated (see Suppl. Figure S1) and as a control synthesized full-length wild-type (WT) *prkar2b*. The two synthesized versions of *prkar2b* were cloned with GFP fused to the N-terminus of PRKAR2B. The level of *prkar2b* transcripts was normalized to *gfp* mRNA to compensate for any differences in transfection efficiency and expression of the 2 plasmids and this showed that ablating the seeds rendered *prkar2b* transcripts resistant to miR-34c-mediated dicing (Fig. 3B, right panel). Taken together, this indicates that *prkar2b* expression is regulated by variations in miR-34c-3p levels and confirms that *prkar2b* mRNA possesses a bona fide miR-34c-3p seed sequence.

**Figure 3:**
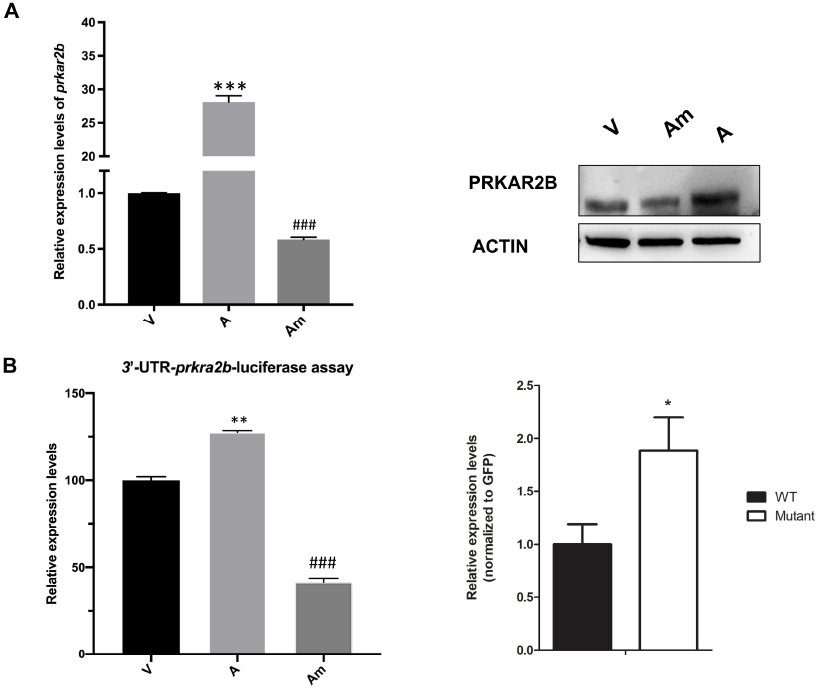
*Prkar2b* is a direct miR-34c-3p-target gene. A. *Theileria*-transformed macrophages transfected with the miR-34c-3p mimic (Am) downregulate both *prkar2b* transcripts (**left panel**) and PRKAR2B protein levels (**right panel)**. **B. Left panel.** Reflecting greater *prkar2b* expression in attenuated macrophages *prkar2b*-luciferase activity is also higher in attenuated (A) than virulent macrophages (V). Luciferase activity decreases when attenuated macrophages are transfected with the miR-34c-3p mimic (Am). **Right panel**. mRNA level of *prkar2b* in virulent macrophages transfected with either WT- or mutant-PRKAR2B expression plasmids. Mutation of the 4 seed sites rendered *prkar2b* transcripts resistant to miR-34c dicing.

We have previously established that dissemination of *T. annulata*-transformed macrophages is cAMP-PKA-dependent (25–27). In Fig.1B we showed that augmenting miR-34c levels increased Matrigel traversal suggesting that ablation of PRKAR2B expression by miR-34c-3p increases PKA activity and macrophage traversal. Virulent macrophages display higher PKA activity compared to attenuated macrophages and transfection of attenuated macrophages with the miR-34c-3p mimic restored PKA activity to virulent levels (Fig. 4, left panel). Thus, by ablating PRKAR2B miR-34c-3p increases both PKA activity and dissemination potential of *Theileria*-transformed macrophages.

**Figure 4:**
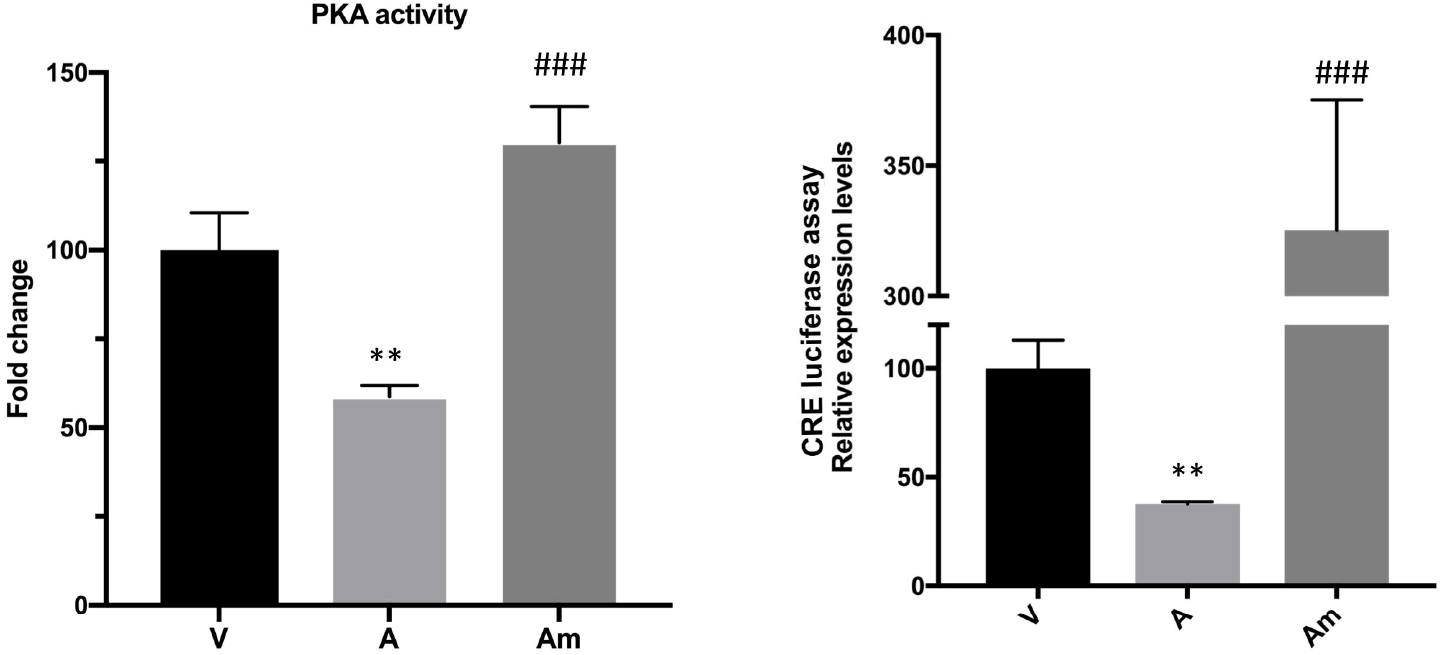
miR-34c-3p-induced drop in PRKAR2B upregulates PKA kinase activity of *T. annulata*-infected macrophages. Left panel. *T. annulata*-infected virulent macrophages (V) have higher PKA activity than attenuated (A) macrophages and overexpression of miR-34c-3p restores PKA activity in attenuated macrophages. **Right panel.** Overexpression of miR-34c-3p in attenuated macrophages increases CREB-transactivation. (Data represented as mean ±SEM. *n*=3. \*\*\**P <* 0.001 compared to V; ###*P <* 0.001 compared to A.

PKA is known to phosphorylate the transcription factor cAMP response element binding protein (CREB) (28,29), so for further confirmation of the capacity of miR-34c to modulate PKA activity the transcription level of CREB was estimated by CRE-driven luciferase assays to show that miR-34c-3p mimic treatment induces an increase in CRE-driven luciferase activity (Fig. 4, right panel). Taken together, this demonstrated that mammalian PKA kinase activity can be regulated independently of changes in cAMP levels via miR-34c-3p-mediated ablation of *prkar2b*.

### miR-34c-3p regulates *P. falciparum*-infected erythrocyte PKA kinase activity

As red blood cell infection by *P. falciparum* has been reported to increase intra-erythrocyte miR-34c-3p levels (see supplementary files in (18)), and given that above we showed that changes in miR-34c-3p levels impacted on PKA activity of *T. annulata*-infected macrophages, we examined if miR-34c-3p can regulate PKA activity of *P. falciparum*-infected RBC. First, we confirmed that infection with *P. falciparum* does indeed lead to increased amounts of intra-erythrocyte miR-34c-3p, and then demonstrated that this increase reduced the abundance of *prkar2b* mRNA and increased PKA activity (Figure 5). Competitive inhibition of miR-34c-3p binding to its cognate seed sequence raised *prkar2b* transcript levels and reduced PKA kinase activity of infected red blood cells (Fig. 5) demonstrating that changes in PKA activity are directly due to miR-34c-3p binding to *prkar2b*.

**Figure 5:**
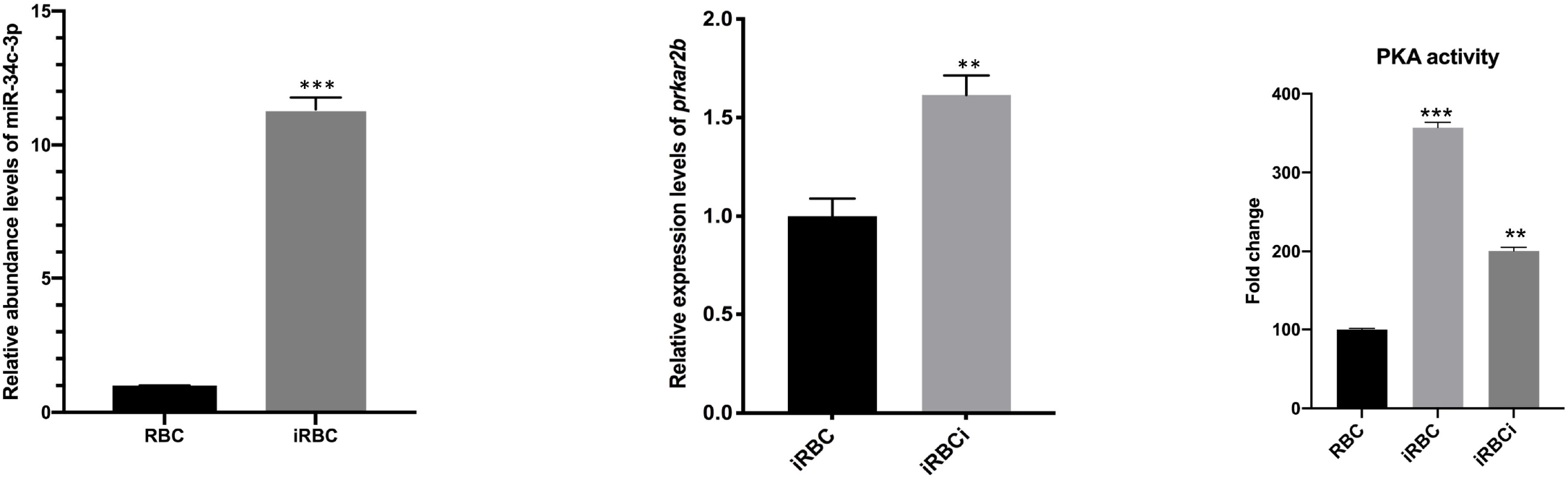
miR-34c-3p targets *prkar2b* to regulate *P. falciparum*-infected erythrocyte PKA activity independently of fluxes in cAMP. Left panel. miR-34c-3p levels drastically increase in infected red blood cells (iRBC) compared to non-infected (RBC). Centre panel. host *prkar2b* is increased in iRBC compared to RBC. Right panel. Infected RBC display higher PKA activity as opposed to non-infected RBC. Inhibition of miR-34c-3p in iRBC (iRBCi) decreases PKA activity. Data represent mean ±SEM. n=3. Data represent mean ±SEM. n=3. **P < 0.005 and ***p < 0.001.

## Discussion

MicroRNA functions have been shown to be possibly associated with neoblast biology, development, physiology, infection and immunity of parasites (31). Importantly, parasite infection can alter host miRNA expression, which can favour both infection and parasite clearance. Here, we characterized the role of miR-34c-3p in modulating the cancer-like phenotype of *Theileria*-infected bovine macrophages. Importantly, we confirmed that *prkar2b* is a direct miR-34c-3p-target gene not only in *T. annulata*-infected macrophages, but also in *P. falciparum*-infected erythrocytes. When the miR-34c-3p seed in the 3’-UTR of *prkar2b* is fused to the luciferase gene the amount of luciferase activity decreases when *T. annulata*-infected macrophages are co-transfected with a miR-34c-3p mimic (Figure 3). Conversely, the ability of miR-34c-3p to ablate infected red blood cell *prkar2b* transcripts was abrogated by treatment with a competitive inhibitor of miR-34c-3p binding (Figure 5).

We demonstrated that miR-34c-3p-induced loss of PRKAR2B enhanced PKA activity to induce CREB transactivation both of which we have shown are important contributors to *Theileria*-transformed macrophage dissemination. We have previously described how TGF-β secreted by *T. annulata*-transformed macrophages contributes to sustaining PKA activity in virulent macrophages via PGE2 engagement of EP4 to augment levels of cAMP (26). Now, we provide an infection-induced epigenetic mechanism for regulating mammalian PKA activity independent of fluxes in cAMP. *T. annulata* infection and transformation of bovine macrophages target mammalian PKA activity in 3 different ways; 1) by raising cAMP levels (26); 2) by suppressing the endogenous inhibitor PKIG (26) and 3) by miR-34c-3p ablation of PRKAR2B subunit expression and this three-pronged “attack” underscores the key role PKA plays in the dissemination of *T. annulata*-transformed macrophages.

In addition, we provide evidence that miR-34c-3p also targets *prkar2b* in *P. falciparum*-infected erythrocytes. First, we confirmed a previous report (18) that miR-34c-3p levels are higher in RBC-infected with *P. falciparum* than in non-infected erythrocytes. Although red blood cell miR-34c-3p targets were not previously characterized (18) we now show that the amount of *prkar2b* mRNA in infected erythrocytes increases upon treatment with an antagonist that competitively inhibits binding to miR-34c-3p seed sequences that increases *prkar2b* abundance and reduces PKA activity (Figure 5). Overall, our results suggest that miR-34c-3p mimics and antagonists could have therapeutic use in the treatment of different diseases caused by intracellular parasites and in Figure 6 we present a generalized schematic representing how miR-34c-3p ablation of PRKAR2B regulates PKA kinase activity independent of fluxes in cAMP in *Theileria*-infected leukocytes and *Plasmodium*-infected erythrocytes.

**Figure 6:**
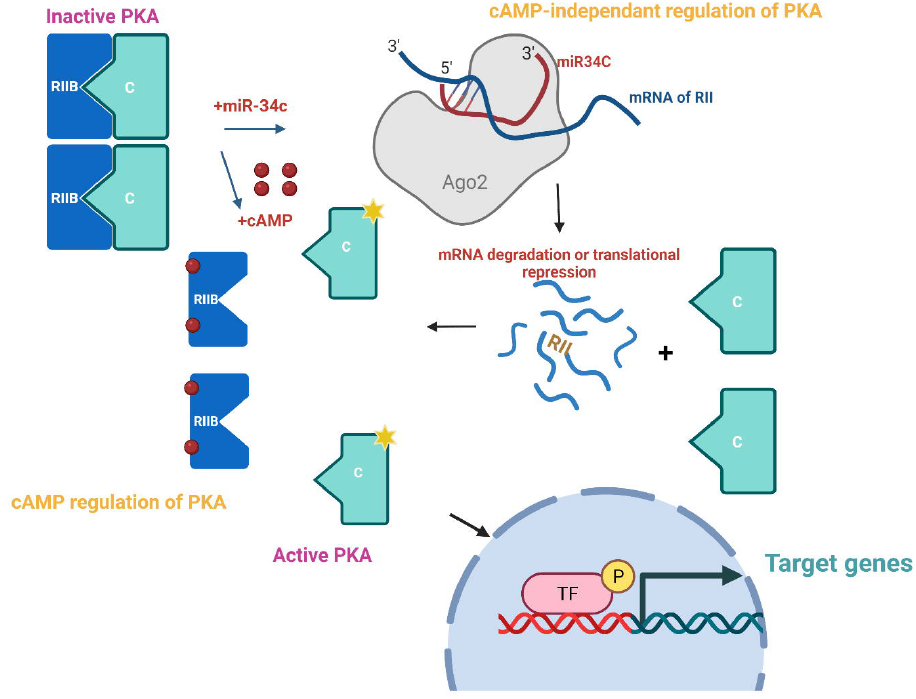
Model proposing how miR-34c-3p by targeting PRKAR2B alters PKA kinase activity independent of fluxes in cAMP. PKA is a heteromer composed of two catalytic subunits bound to two regulatory subunits. Classical activation of PKA occurs when cAMP binds to regulatory subunits causing a conformational change in catalytic subunits from an inactive to an active kinase. miR-34c-3p can regulate PKA catalytic activity independently of cAMP by ablating *prkar2b* leading to reduced PRKAR2B expression and greater PKA catalytic activity towards target proteins in the nucleus and cytosol.

## Materials and Methods

### Cell culture

Cells used in this study are *T. annulata*-transformed Ode macrophages, where virulent macrophages used corresponds to passage 62 and attenuated macrophages to passage 364 (32). Non-infected BL20 and *T. annulata*-infected BL20 (TBL20) cells used have been previously characterized (33, 34). All cells were incubated at 37°C with 5% CO_2_ in Roswell Park Memorial Institute medium (RPMI) supplemented with 10% Fetal Bovine Serum (FBS), 2 mM L-Glutamine, 100 U penicillin, 0.1 mg/ml streptomycin, and 4-(2-hydroxyethyl)-1-piperazineethanesulfonic acid (HEPES) and 5% 2-mercaptoethanol for BL20 and TBL20.

### Malaria parasite culturing

*P. falciparum* 3D7 were cultured in RPMI 1640 (Life Technologies, 51800-035) supplemented with 0.5% (w/v) Albumax II (11021-29), 200 μM hypoxanthine, 20 μg/mL gentamycin (complete RPMI 1640 medium) in human erythrocytes at a hematocrit between 2.5% and 5%. Parasite cultures were maintained under low oxygen pressure (1% Ob_2_, 3% CO_2_, 96% N_2_) at 37°C.

### RNA extractions

Total RNA of *Theileria*-infected leukocytes was isolated using the Direct-zol RNA Kit (Zymo Research # R2070) and total RNA designated for miRNA experiments was extracted using the mirVana miRNA isolation kit (Thermo Fisher) according to the manufacturer’s instructions. The quality of extracted RNA was verified using a Bioanalyzer 2100 and quantification was carried out using Qubit (Invitrogen, catalogue number Q10210).

### qRT-PCR for miRNAs

RNA extracted with the mirVana kit was used. cDNA was synthesized using the TaqMan microRNA RT kit (Applied Biosystems) following the manufacturer’s instructions. A total of 10 ng of RNA was used in the reverse transcription reaction with miRNA-specific primers. The real-time reactions were performed in a 7500 HT Fast Real-time PCR system (Applied Biosystems). Data were analysed using the 2^−ΔΔCT^. The cycle threshold (Ct) values from the selected miRNA targets were subtracted from the Ct values of the endogenous small noncoding RNA control RNU6B (control miRNA assay; Applied Biosystems).

### qRT-PCR for mRNA

Total RNA was reverse transcribed using the High-Capacity cDNA Reverse Transcription Kit (Applied Biosystems, catalogue number 4368814) as follows: 1 *μ* g of total RNA, 2 *μ* L of RT buffer, 0.8 *μ* L of 100mM dNTP mix, 2.0 *μ* L of 10X random primers, 1 *μ* L of MultiScribe reverse transcriptase and Nuclease-free water to a final volume of 20 *μ* L.

The reaction was incubated for 10 min at 25°C, 2 h at 37°C then the enzyme was inactivated at 85°C for 5 min. Real-time PCR was performed in a 20μL reaction containing cDNA template, 10 *μ* L 2X Fast SYBR Green Master Mix and 500 nM of forward and reverse primers. The reaction was run on the 7500 HT Fast Real-Time PCR System (Applied Biosystems). Expression of *gapdh, hprt1* and *beta-actin* were used as measures of housekeeping gene expression and the results were analyzed by the 2^−ΔΔCT^ method. The error bars represent the SEM of 3 biological replicates.

### Western Blotting

Cells were harvested and extracted by 1x mammalian lysis buffer (Abcam # ab179835) supplemented with protease and phosphatase inhibitor cocktail (#78440, Thermo Fisher). Protein concentration was determined by the Bradford protein assay. Cell lysates were subjected to Western blot analysis using conventional SDS/PAGE and protein transfer to nitrocellulose filters (Protran, Whatman). The membrane was blocked by 5% non-fat milk-TBST (for anti-PRKAR2B and anti-GAPDH), or 3% non-fat milk-PBST (for anti-actin antibody) overnight at 4 ^0^C. Antibodies used in immunoblotting were as follows: rabbit polyclonal antibody anti-GAPDH (Merck Millipore #ABS16), mouse monoclonal antibody anti-PRKAR2B (#610625, BD transduction laboratories) and goat polyclonal antibody anti-actin (Santa Cruz Biotechnologies I-19). After washing, proteins were visualized with ECL western blotting detection reagents (Thermo Scientific) with a fusion instrument. The β-actin was used as a loading control throughout all experiments.

### Fluorescent activated cell sorting (FACS) of GFP-PRKAR2B expressing cells

Wild type *prkar2b* subunit of bovine PKA and mutant *prkar2b* (harbouring mutations that ablate all potential miR-34c seed sites) were synthesized (GenScript) and cloned into pmaxGFP vector (LONZA). Virulent *T. annulata*-transformed macrophages were transfected with plasmids expressing wild-type- or mutant-PRKAR2B with GFP fused to the N-termini. The transfection rate (efficiency) was measured, and in 3-independent transfections, the efficiency averaged 19%. Therefore, to obtain a high percentage of transfected cells 24 h post-transfection GFP expressing cells were FACS sorted by MoFlo Astrios (Beckman). Following sorting total RNA was prepared, as described above.

### Matrigel Chamber assay

The dissemination capacity of Ode macrophages was assessed *in vitro* using Matrigel migration chambers, as described (35). The Culture Coat Medium BME (basement membrane extract) 96-well assay was performed according to Culturex instructions (catalog number 3482-096-K). After 24 h of incubation at 37°C, each well of the top chamber was washed once in the buffer. The top chamber was placed back on the receiver plate. One hundred microliters of cell dissociation solution-Calcein AM was added to the bottom chamber of each well, and the mixtures were incubated at 37°C for 1 h with fluorescently labelled cells to dissociate the cells from the membrane before reading at 485-nm excitation and 520-nm emission wavelengths, using the same parameters as those used for the standard curve.

### Transfection

Macrophages were transfected by electroporation using the Nucleofector system (Amaxa Biosystems). A total of 5 × 10^5^ cells was suspended in 100 μL of Nucleofector solution mix with 400 pM of mimic of miR-34c-3p and subjected to electroporation using the cell line-specific program T-O17 for *Theileria*-infected macrophages. After transfection cells were suspended in a fresh complete medium and incubated at 37°C with 5% CO_2_ for 48 h.

### Synchronization of *P. falciparum*-infected red blood cell culture and pharmacological inhibition or stimulation of miR-34c-3p

Parasites were cultured to at least 10% parasitemia in T-75 flasks containing 25 mL medium at 2% hematocrit. Parasites were synchronized first at ring stage with two rounds of 5% sorbitol. When most of the parasites had matured to schizonts, the culture was loaded on 70% Percoll cushion and centrifuged for 10 min at 800x g. Schizonts from the top of 70% Percoll were collected washed two times with complete media and re-suspended in complete medium. Parasitemia was adjusted to 0.2-0.5% and 2% hematocrit and the culture was distributed in triplicates in 6-wells plate with various treatments: 400pM of the mimic (HMI0513-5NMOL, Sigma) or 400pM of the inhibitor (cat# 4464084, ID: MH12342, Ambion from Thermo fisher). The culture was incubated at 37°C with 5% CO_2_/3% O_2_/balanced N_2_ gas mixture, for 96 h. To follow growth, blood smears were prepared and stained with Giemsa solution (Sigma).

### CREB luciferase assay

After transfection, cells were suspended in fresh complete medium and incubated at 37°C, 5% CO_2_ for 24 h and cells were lysed after 48 h. Measurements of luciferase and ß-galactosidase activities were performed using the Dual Light Assay system (Life Technologies) and luminometer Centro LB 960 (Berthold) according to the manufacturer’s instructions.

### Measurement of PKA activity

*Theileria*-infected macrophages were transfected with the specific mimic of miR-34c-3p. Samples were then collected, centrifuged at 1500 rpm for 5 min and lysed by recommended lysis buffer [20 mM MOPS, 50mM β-glycerolphosphate, 50mM sodium fluoride, 1 mM sodium vanadate, 5 mM EGTA, 2 mM EDTA, 1% NP40, 1mM dithiothreitol (DTT), 1 mM benzamidine, 1mM phenylmethane-sulphonylfluoride (PMSF) and 10 μg/mL leupeptin and aprotinin. For permeabilization of *P. falciparum*-infected red blood cells, Streptolysin O (SLO, Sigma) was first titrated by re-suspending SLO (25000 units) in 5 mL of PBS (10 mM sodium phosphate buffer and 145mM NaCl, pH 7,4) containing 0,1 % BSA and activating by incubation with 5-10 mM dithiothreitol at 37°C for 2 h. iRBCs were re-suspended in 200 μL of RPMI containing 3-4 hemolytic units of SLO, resulting in haemoglobin release of more than 98%. Following permeabilization with SLO, cells were centrifuged and washed with 200 μL of RPMI. *P. falciparum*-infected erythrocytes pellets were lysed by the recommended buffer and lysates were cleared by centrifugation (15,000 rpm for 15 min at 4°C), and the total amount of proteins in the supernatant was measured by a Bio-Rad protein assay, based on the method of Bradford, using BSA as a standard. PKA activity was measured using an ELISA kit (PKA kinase activity kit, Abcam 139435) according to the manufacturer’s instructions.

### Statistical Analysis

Data were analysed with the Student’s two-tailed T-tests with 3 biological replicates for each experiment. All values are expressed as mean+/-SEM. In each figure, the following symbols represent respective p-value ranges: *:P<0.05; **: p<0.005, ***: p < 0.001.

## Acknowledgments

This study was supported by a Competitive Research Grant (CRG) from the Office for Sponsored Research (OSR-2015-CRG4-2610) at King Abdullah University of Science and Technology (KAUST) awarded to AP and GL. MH, FBR and ZR acknowledge KAUST for Postdoctoral fellowships. SM and TB were supported by AP baseline funding (BAS/1/1020-01-01) from KAUST. GL also acknowledges ANR-11-LABX-0024 and core support from INSERM and the CNRS. FACS was performed with the help of the cytometry and immuno-biology facility (CYBIO) of the Cochin Institute – INSERM U1016.

## Supporting information

**Figure S1:** Sequences of wild-type (WT) and mutant *prkar2b* with all 4 predicted non-canonical miR-34 seeds in the coding sequence mutated. Original seed sites are highlighted in yellow and mutated sites in green.

