## Supplemental figure 1 for "miR-34c-3p regulates PKA activity independent of cAMP via ablation of PRKAR2B in *Theileria annulata*-infected leukocytes"

**PRKAR2B-WT**

atg agc atc gag atc cca gcg gga ctg acg gag ctg ctg cag ggc ttc acc gtg gag gtc
M S I E I P A G L T E L L Q G F T V E V
ctg agg cac cag ccc gcg gac ctg ctg gag ttc gcg ctg cag cac ttc aca cgc ctg cag
L R H Q P A D L L E F A L Q H F T R L Q
gag gag aac gag cgc aaa ggc acc gcg cgc ttc ggc cat gaa ggc agg acc tgg ggg gac
E E N E R K G T A R F G H E G R T W G D
gcg ggc gcc gcc ggc ggg ggc ggc acc ccc agc aag ggg gtc aac ttc gcc gag gag ccc
A G A A G G G G T P S K G V N F A E E P
agg cac agc gat tcg gag aac ggc gag gag gag gag gaa gag gcc gcg gat gca ggg gca
R H S D S E N G E E E E E E A A D A G A
ttc aac gct cca gta ata aac cga ttt aca agg cgt gcc tca gta tgt gca gaa gct tat
F N A P V I N R F T R R A S V C A E A Y
aat cct gat gaa gaa gaa gat gat gct gaa tcc agg att ata cat ccc aaa act gac gat
N P D E E E D D A E S R I I H P K T D D
caa aga aat cga ttg caa gag gct tgc aaa gac atc ctg cta ttt aag aac ctg gat ccg
Q R N R L Q E A C K D I L L F K N L D P
gag cag atg tct caa gta tta gat gcc atg ttt gaa aaa ttg gtt aaa gaa ggg gaa cat
E Q M S Q V L D A M F E K L V K E G E H
gta att gat caa ggt gat gat ggc gac aac ttt tat gta att gat aga gga acg ttt gat
V I D Q G D D G D N F Y V I D R G T F D
att tat gtg aaa tgc gat ggt gtt gga cgg tgt gtt ggc aac tat gac aat cgt ggg agt
I Y V K C D G V G R C V G N Y D N R G S
ttt ggc gaa ctg gcc tta atg tac aac aca ccc aga gca gct aca atc act gct acc tct
F G E L A L M Y N T P R A A T I T A T S
cct ggt gct ctg tgg ggt ttg gac agg gta acc ttc agg agg ata att gta aaa aat aat
P G A L W G L D R V T F R R I I V K N N
gcc aaa aag aga aaa atg tat gaa agc ttt att gag tca ctg cca ttc ctt aaa tct ttg
A K K R K M Y E S F I E S L P F L K S L
gaa gtt tct gaa cgc ctg aaa gtg gta gac gtg ata ggc acc aaa gta tac aac gat gga
E V S E R L K V V D V I G T K V Y N D G
gaa caa atc att gct cag gga gat tcg gct gat tct ttt ttc att gta gaa tcg gga gaa
E Q I I A Q G D S A D S F F I V E S G E
gtg aaa att act atg aaa agg aag ggt aag tca gag gtg gag gag aac ggt gca gtt gaa
V K I T M K R K G K S E V E E N G A V E
atc gct cgg tgt tct cga ggg cag tac ttc ggc gaa ctg gct ctg gta acg aac aag ccc
I A R C S R G Q Y F G E L A L V T N K P
cga gcc gct tcc gca cac gcc atc ggg acc gtc aaa tgt tta gcc atg gat gtg caa gca
R A A S A H A I G T V K C L A M D V Q A
ttt gaa agg ctt ttg gga cct tgc atg gaa att atg aaa agg aac atc gct acc tat gaa
F E R L L G P C M E I M K R N I A T Y E
gag caa tta gtt gcc ctg ttt gga aca aac atg gat att gtt gaa ccc act gca tga

**PRKAR2B-Mutant**

atg agc atc gag atc cca gcg gga ctg acg gag ctg ctg cag ggc ttc acc gtg gag gtc
M S I E I P A G L T E L L Q G F T V E V
ctg agg cac cag ccc gcg gac ctg ctg gag ttc gcg ctg cag cac ttc aca cgc ctg cag
L R H Q P A D L L E F A L Q H F T R L Q
gag gag aac gag cgc aaa ggc acc gcg cgc ttc ggc cat gaa ggc agg acc tgg ggg gac
E E N E R K G T A R F G H E G R T W G D
gcg ggc gcc gcc ggc ggg ggc ggc acc ccc agc aag ggg gtc aac ttc gcc gag gag ccc
A G A A G G G G T P S K G V N F A E E P
agg cac agc gat tcg gag aac ggc gag gag gag gag gaa gag gcc gcg gat gca ggg gca
R H S D S E N G E E E E E E A A D A G A
ttc aac gct cca gta ata aac cga ttt aca agg cgt gcc tca gta tgt gca gaa gct tat
F N A P V I N R F T R R A S V C A E A Y
aat cct gat gaa gaa gaa gat gat gct gaa tcc agg att ata cat ccc aaa act gac gat
N P D E E E D D A E S R I I H P K T D D
caa aga aat cga ttg caa gag gct tgc aaa gac atc ctg cta ttt aag aac ctg gat ccg
Q R N R L Q E A C K D I L L F K N L D P
gag cag atg tct caa gta tta gat gcc atg ttt gaa aaa ttg gtt aaa gaa ggg gaa cat
E Q M S Q V L D A M F E K L V K E G E H
gta att gat caa ggt gat gat ggc gac aac ttt tat gta att gat aga gga acg ttt gat
V I D Q G D D G D N F Y V I D R G T F D
att tat gtg aaa tgc gat ggt gtt gga cgg tgt gtt ggc aac tat gac aat cgt ggg agt
I Y V K C D G V G R C V G N Y D N R G S
ttt ggc gaa ctg gcc tta atg tac aac aca ccc aga gca gct aca ata acg gct acc tct

F G E L A L M Y N T P R A A T I T A T S

cct ggt gct ctg tgg ggt ttg gac agg gta acc ttc agg agg ata att gta aaa aac aat

P G A L W G L D R V T F R R I I V K N N

gca aaa aag aga aaa atg tat gaa agc ttt att gag tcc tta ccg ttc ctt aaa tct ttg

A K K R K M Y E S F I E S L P F L K S L

gaa gtt tct gaa cgc ctg aaa gtg gta gac gtg ata ggc acc aaa gta tac aac gat gga

E V S E R L K V V D V I G T K V Y N D G

gaa caa atc att gct cag gga gat tcg gct gat tct ttt ttc att gta gaa tcg gga gaa

E Q I I A Q G D S A D S F F I V E S G E

gtg aaa att act atg aaa agg aag ggt aag tca gag gtg gag gag aac ggt gca gtt gaa

V K I T M K R K G K S E V E E N G A V E

atc gct cgg tgt tct cga ggg cag tac ttc ggc gaa ctg gct ctg gta acg aac aag ccc

I A R C S R G Q Y F G E L A L V T N K P

cga gcc gct tcc gca cac gcc atc ggg acc gtc aaa tgt tta gcc atg gat gtg caa gca

R A A S A H A I G T V K C L A M D V Q A

ttt gaa agg ctt ttg gga cct tgc atg gaa att atg aaa agg aac atc gct acc tat gaa

F E R L L G P C M E I M K R N I A T Y E

gag caa tta gtt gcc ctg ttt gga aca aac atg gat att gtt gaa ccg acg gca tga
